# Mucosally sourced complement factor B modulates the host response to colitis

**DOI:** 10.64898/2025.12.31.697141

**Authors:** Aayusha Thapa, Aasritha Nallapu, Nnenna Mougboh, Swathi Nedunchezian, Khushi Talati, Brian Yang, Jungheun Hyun, John Michael Sanchez, Christopher S. Seet, Hon Wai Koon, Zhaoping Li, Matthew A. Ciorba, Kathrin Michelsen, Hrishikesh. S. Kulkarni, Devesha H. Kulkarni

## Abstract

Distinct host factors maintain intestinal homeostasis but are incompletely understood. The complement system is primarily liver-derived and serum-operative. However, there is growing recognition for complement-mediated host defense at mucosal surfaces. The alternative pathway, which is constitutively active at low levels and amplifies complement activation independent of antibodies, requires Complement Factor B (CFB). Despite its evolutionary conservation, the spatial, cellular, and functional roles of CFB in the intestine are poorly understood. Here, we show that CFB is produced in the human colon and is increased in patients with active inflammatory bowel disease. To isolate the role of local CFB in mucosal responses, we interrogated a mouse strain that has no circulating, liver-derived CFB but retains intact CFB expression in the gut. Global CFB-deficient mice succumb to colitis compared to these liver-specific knockout mice, suggesting that locally synthesized CFB mitigates colitis. Single-cell analyses identify enterocytes and fibroblasts as key CFB producers in the gut. Compartment-specific deletion of CFB from epithelial or stromal cells abrogates mucosal protection independent of circulating levels, which corroborates with pharmacological CFB inhibition. These findings redefine complement in the intestine as a locally regulated mucosal defense system and establish gut-derived CFB as a critical determinant of intestinal homeostasis.

**BRIEF SUMMARY:** The role of local immune mediators in gut mucosal immune responses is still not entirely understood. In this study, we demonstrate a novel role for complement protein Factor B, a key component of the alternative pathway, which is locally sourced through epithelial and stromal cells. In vivo modeling of impaired local Factor B synthesis results in worse colitis, revealing a key role for mucosal sourced components of the alternative pathway.

## INTRODUCTION

The intestinal barrier is comprised of highly specialized cells that maintains homeostasis, which when altered, results in chronic inflammation^1^. Hence, it is necessary to understand the distinct host factors that contribute to the maintenance of intestinal homeostasis. Dysregulated intestinal immune homeostasis can result in inflammatory bowel disease (IBD), which affects nearly 3 million adults in the United States, and its prevalence continues to rise^2, 3^. The current treatment strategies for IBD include immunomodulators and biologic agents such as anti-TNF therapies. However, these medications can result in significant side effects, and 1 in 5 patients become refractory to medical therapy^4^. Additionally, most of these medications do not facilitate wound healing or target the mucosal barrier. Hence, there is a critical need for new treatment strategies, including those that promote mucosal homeostasis and maintain host defense.

The complement system is an evolutionarily conserved family of proteins that facilitate immune responses by attacking microbes to decrease pathogen burden^5, 6^. Genetic complement deficiencies are associated not only with recurrent bacterial infections^6^ but also with severe complications of IBD^7, 8, 9^. Traditionally, the effects of most complement proteins were attributed to their origins in the liver and functions in the fluid-phase (i.e., circulation^10^). However, we and others have recently demonstrated the importance of local synthesis of C3 (a central complement component) at mucosal sites such as the lung and the kidney^11, 12, 13^. C3 is also expressed in colonic stromal cells, and enteric infection stimulates local C3 levels^12^. However, few, if any, studies have interrogated how local complement activity modulates gut mucosal homeostasis and how Factor B (**CFB**, human gene: *CFB*, mouse gene: *Cfb)*, a key component of the alternative pathway (AP) of complement activation, is relevant to mucosal health.

Although complement has been implicated in IBD since 1970s^14, 15^, it is unclear as to how it affects disease pathogenesis. GWAS studies have revealed that a functional polymorphism in *CFB* (*rs4151651,* minor allele frequency: 2-5%) is strongly associated with perianal Crohn’s disease and medically refractory ulcerative colitis^7, 8, 9^. *rs4151651* leads to a substitution from glycine to serine in *CFB*, resulting in impaired binding of Factor B to activated C3 fragments (i.e., C3b) and impaired cleavage of Factor B into its active Ba and Bb subunits^7^. This SNP thus impairs AP activation, which has led to speculation that its effects in IBD are due to increased bacterial burden in the setting of decreased phagocytosis^7^. Additionally, the sourcing of individual complement proteins at the site of injury, and how a balance is maintained between an adequate gut mucosal host response versus the dysregulation observed in IBD remains to be answered. For example, although impaired AP activation is associated with IBD complications, increased C3 and Factor B have also been observed in IBD^14, 15, 16, 17^. *CFB* mRNA is elevated in inflamed versus normal colonic mucosal biopsies in patients with IBD^18^, suggesting local Factor B production may be contributing to uncontrolled complement activation and tissue damage. Hence, understanding how the AP is sourced, activated and regulated in colitis, and how it can be modulated locally to prevent tissue damage will facilitate developing targeted, host-focused therapies in IBD.

Given its strong genetic association with IBD, we sought to address the role of Factor B in intestinal homeostasis and function. We demonstrate the extrahepatic sourcing of Factor B in the colon and the significance of gut-derived Factor B in the context of mucosal immunity. Using single cell and spatial transcriptomic analyses, and histological comparisons of human IBD specimens, we demonstrate a strong correlation between increased local activation of Factor B and active disease. Using a murine model of acute colitis and in vitro systems, we further demonstrate a critical role of Factor B derived from structural cells forming the mucosal barrier, notably epithelial cells and fibroblasts. Taken together, our study demonstrates that mucosal sourced Factor B increases during acute intestinal inflammation but also plays a critical role mitigating colitis severity, and thus, protects the barrier.

## RESULTS

### Factor B is upregulated in inflamed colons during IBD

Since complement proteins have recently been reported at various mucosal surfaces including the gut^12^, we assessed complement gene expression in patients with IBD. We queried single cell RNA sequencing (scRNA-seq) data published using uninflamed and inflamed colon tissue from 18 patients with ulcerative colitis (UC) under different treatment regimens and 12 healthy non-IBD individuals^19^. We observed that alternative pathway (AP) components, notably *CFB,* are increased in the inflamed tissues compared to that in the non-inflamed and healthy tissues (**Fig. 1A**). *CFB* expression was significantly higher in the inflamed tissue compared to healthy specimens amongst epithelial, immune and stromal cell subsets based on the cluster identification (**Fig. 1B, Supplementary Fig. S1A**). Further, we interrogated other AP components such as *C3*, Factor D (*CFD*) and properdin (*CFP*), and observed that upregulation in these other components were not as pronounced across the type of tissue and cell types (**Supplementary Fig. S1B-D).** To validate our observations in other human cohorts with colitis, we assessed complement gene expression in samples of patients with checkpoint-inhibitor colitis^20^. We observed that *CFB* expression was increased in inflamed tissues compared to that in the non-inflamed and healthy tissues (**Supplementary Fig. S1E**) with a significantly higher expression in epithelial and stromal cell subsets (**Supplementary Fig. S1F**).

**Figure 1.**
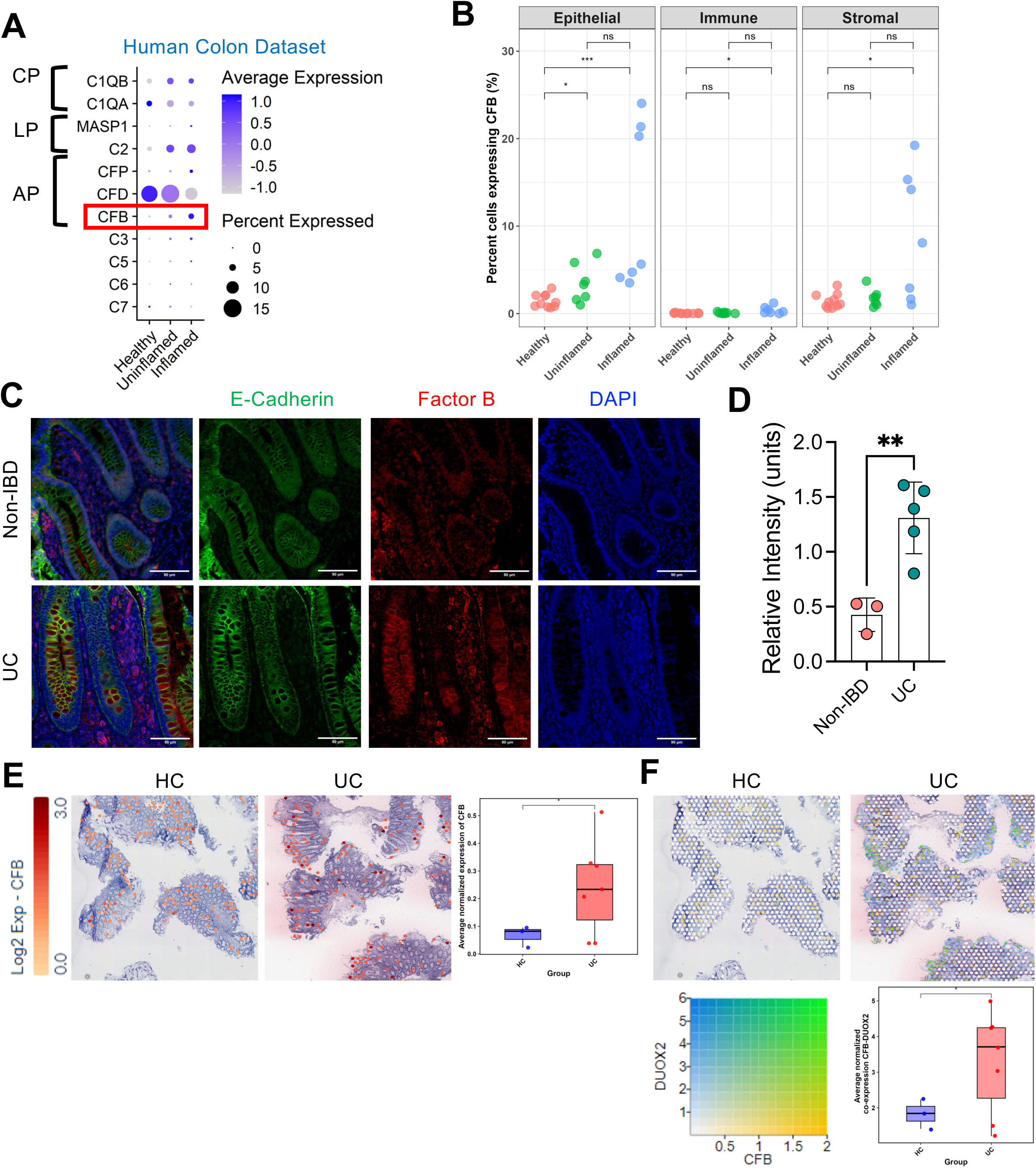
Increased intestinal Factor B expression in patients with IBD. (**A**) Dot plot illustrating the expression of complement genes representing the classical (CP), lectin (LP) and alternative (AP) pathway in intestinal tissue biopsies from uninflamed and non-inflamed regions of patients with ulcerative colitis (UC, Single Cell Portal: SCP259). (**B**) Plots depict *CFB* expression across denoted colonic cell types in the same dataset from intestinal tissue of patients with UC and healthy controls. * P <0.05, *** P <0.001, ns, not significant, Wilcoxon rank test. (**C**) Immunofluorescence staining for Factor B (red), E-cadherin (green) and DAPI (blue) in colonic tissue specimens from patients with IBD versus controls (non-IBD). (Scale bar = 90*μ*m). (**D**) Quantification of the relative intensity of Factor B expression across tissue type. (**E**) Representative spatial transcriptomic slides (Visium) showing *CFB* expression in intestinal tissues obtained from patients with UC compared to healthy controls (HC, GSE189184). Graph depicts quantification of *CFB* expression from 3 HC and 7 UC specimens. (**F**) Representative spatial transcriptomic slides showing DUOX2-CFB co-localization spots in HC (left) and UC tissue (right) from the same dataset. Bar plot represents average co-expression across tissue samples. Error bars represent standard error of the mean (SEM). * P < 0.05, unpaired t-test.

To interrogate whether the proteins levels of Factor B align with its gene expression in the gut, we compared its staining in colon biopsy specimens from inflamed tissue of UC patients (n=5) compared to those who did not have IBD (n=3) (**Fig. 1C**). The intensity of Factor B staining was significantly higher in the inflamed gut tissue from patients with UC (**Fig. 1D**). Further, to dissect the spatiotemporal expression of key AP components in intestinal tissues of patients with IBD, we interrogated *CFB* and *C3* expression in a spatial transcriptomics (Visium) dataset comparing inflamed UC biopsy specimens to non-IBD specimens^20^. *CFB* expression was higher in the inflamed UC biopsies (**Fig. 1E**), whereas C3 expression remained largely comparable between the groups (**Supplementary Fig. S1G**). *CFB* expression appeared predominantly in the epithelial compartment (**Fig. 1E**). Given that DUOX2 (Dual oxidase 2), a hydrogen-peroxide generator, is often used as a marker for early-stage IBD and is expressed by epithelial cells^21^, we assessed the co-expression of *CFB* and *DUOX2* in this dataset. We observed increased co-expression of *CFB* and *DUOX2* in inflamed UC tissues compared to the non-IBD tissue specimens (**Fig. 1F**). Taken together, this data indicates that Factor B is increased in inflamed regions in UC and is likely being sourced from the epithelial barrier.

### Deletion of hepatic-sourced Factor B uncovers role of gut-derived Factor B

To investigate the functional relevance of Factor B made in the colon, we developed a new *Cfb*^f/f^*AlbCre^+/ -^*mouse (mice lacking liver-derived Factor B) and compared it to a newly generated whole body Factor B-deficient mouse (*Cfb^−/−^*, sequence **Supplementary Fig. S2A**). These mice have no circulating Factor B (**Fig. 2A**) yet produce it in the gut at levels comparable to Factor B-sufficient littermate mice (*Cfb^f/f^*, **Fig. 2B-D**, sequence **Supplementary Fig. S2B**). To test if the absence of Factor B alters the gut epithelial barrier, we measured soluble CD14 in plasma from our mice and observed that there was no change in the levels of circulating sCD14, unless the mice were injured using lipopolysaccharide (LPS, **Fig. 2E**). Additionally, to test if gut microbes alter the expression of *Cfb* under homeostasis, we compared *Cfb* expression in colonic epithelial cells from germ-free as well as specific pathogen free (SPF) mice. Our data suggest no difference in *Cfb* expression between germ-free and SPF mice (**Supplementary Fig. S2C**). To identify which intestinal cell types synthesize Factor B, we performed scRNA-Seq on healthy colonic tissue from non-injured *Cfb^f/f^*(i.e. *Cfb^f/f^AlbCre^−/−^*) mice. Supporting our observations from the human cohorts, we observed that *Cfb* is predominantly expressed in the intestinal epithelial cell, stromal cell, and, to a lesser extent, in the immune cell cluster in the gut (**Fig. 2F**). Taken together, we conclude that the colon produces Factor B independent of its circulating levels.

**Figure 2.**
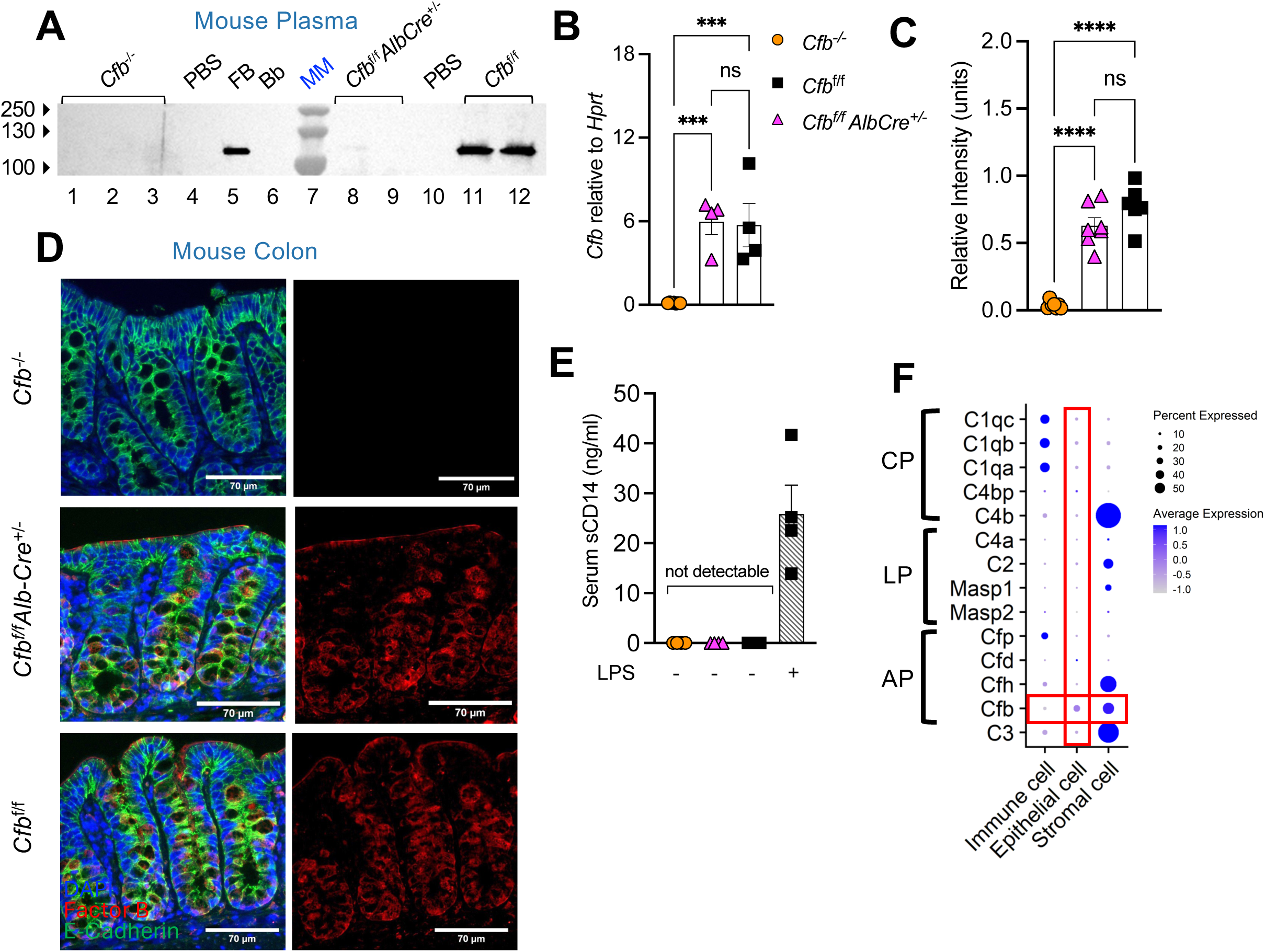
Extrahepatic production of Factor B in the mouse intestine. (**A**) Immunoblot denoting Factor B (FB) in plasma of global Cfb knockouts (*Cfb^−/−^*), mice deficient in liver-derived FB (*Cfb*^f/f^*AlbCre*^+/−^) and their littermates (*Cfb*^f/f^*AlbCre*^−/−^). MM: molecular marker. The same volume of plasma was loaded per well at 1:20 dilution. (**B**) *Cfb* expression on qRT-PCR in the colons from indicated mice, values normalized to *Hprt1*. (**C**) Quantification of the relative intensity of Factor B expression in colon tissue from indicated mice. (**D**) Representative immunofluorescence images of colonic tissue from *Cfb^−/−^*, *Cfb*^f/f^*AlbCre*^+/−^ and *Cfb*^f/*f*^*AlbCre^−/−^* mice (n=4/group), stained with Factor B (red), E-cadherin (green) and DAPI (blue). Scale bars = 70*μ*m. (**E**) Graph depicts serum CD14 concentration. Serum from mice challenged with lipopolysaccharide (LPS) were used as a positive control for barrier leak. Each dot represents an individual mouse. Bar graphs show the mean ± SEM, * P < 0.05; ** P < 0.01, ns, non-significant, unpaired *t* test. (**F**) Dot plot illustrating the expression of complement genes representing the classical (CP), lectin (LP) and alternative (AP) pathway across cell types indicated along the Y axis, based on single-cell RNA-seq data from the large intestines of non-injured adult C57BL/6 mice (*Cfb^f/f^*), housed in specific pathogen-free conditions. Cells pooled from n=2 mice.

### Cytokine stimulation results in Factor B production from the gut

To identify if gut-derived Factor B is increased over time during inflammation, we tested its production by three cell types that partake in innate immune responses in the gut – epithelial, stromal and myeloid cells. To assess the epithelial cell response, we used Caco-2 cells, a human colonic adenocarcinoma cell line. We observed that Caco-2 cells secrete Factor B into the culture supernatant at baseline, and this secretion is significantly increased upon treatment with IL-1β and TNF-α (**Fig. 3A**). We also measured C3 expression in supernatant of the epithelial cells, which paralleled Factor B secretion (**Supplementary Fig. 3A**).

**Figure 3.**
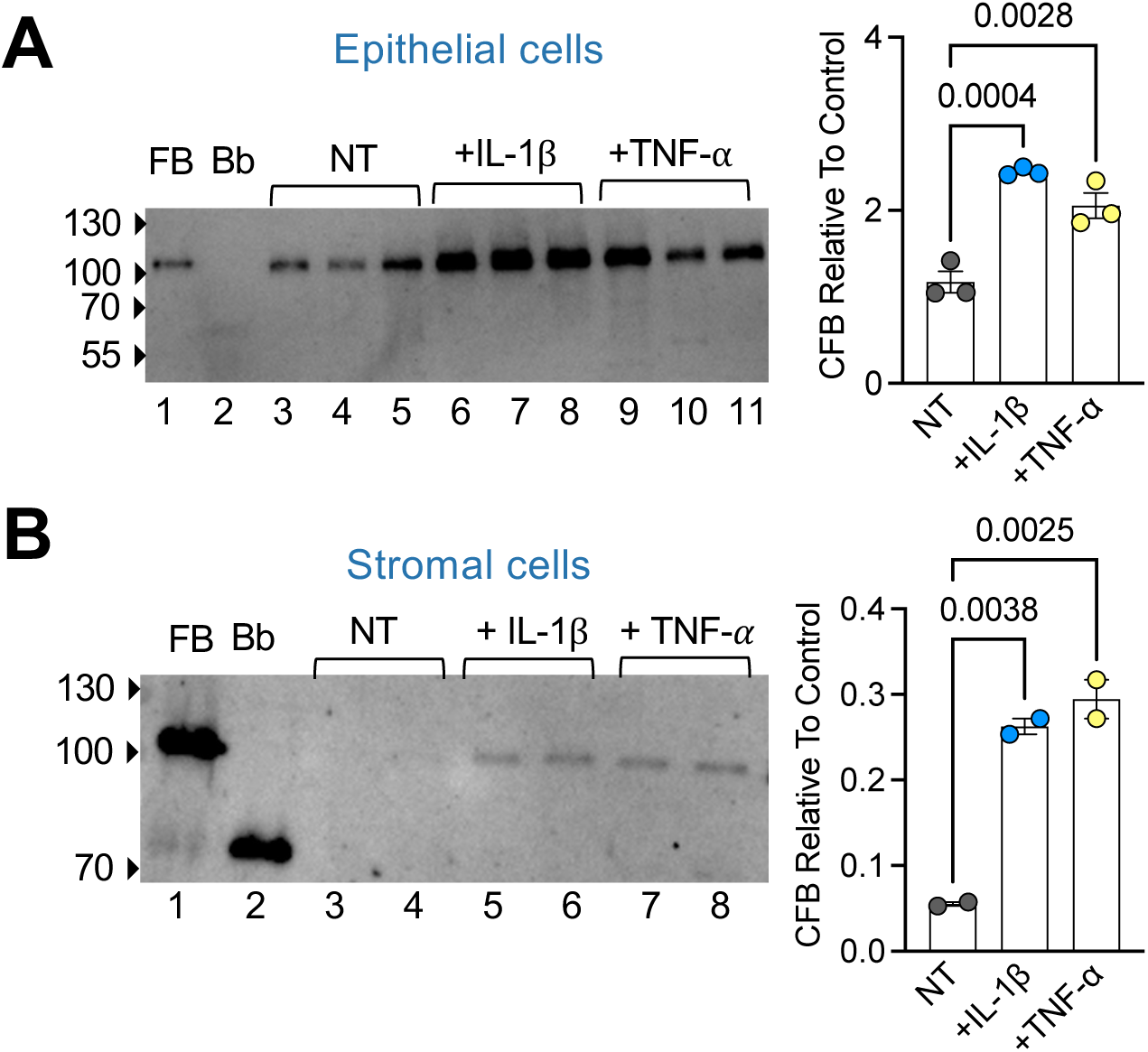
Cytokine-mediated increase of Factor B secretion from colonic cells following stimulation with IBD-relevant cytokines. (**A-B**) Human epithelial Caco-2 cells were incubated in serum-free media alone (non-treated, NT), media with rIL-1β (50ng/ml) or TNF-α (50ng/ml). (**A**) Supernatant was collected after 24 hours, concentrated 50-fold and analyzed by immunoblotting for Factor B. (**B**) Human primary fibroblasts were incubated in serum-free media alone (NT), media with rIL-1β (50ng/ml) or TNF-α (50ng/ml). Immunoblot depicts CFB in supernatant at the 24 h time point. Graph depicts Factor B protein quantification relative to the loaded control. The same volume of supernatant was loaded in each well. Each dot represents an individual replicate. Bar graphs show the mean ± SEM. All immunoblots were repeated at least twice. Statistics performed using an ordinary one-way ANOVA test adjusted for multiple comparisons.

As our transcriptomic analyses reveal that both mouse and human stromal cells express *CFB* (**Fig. 1A & 2E**), we tested whether treatment with IL-1β and TNF-α affects Factor B sourced from cultured primary fibroblasts. We observed a significant increase in Factor B secretion by human colonic fibroblasts post-cytokine stimulation (**Fig. 3B)**. As colonic fibroblasts are known to express C3^12^, we confirmed that treatment with IL-1β and TNF-α also increases C3 levels in their culture supernatant (**Supplementary Fig. S3B**). Finally, we also interrogated the human monocytic cell line THP-1, given that myeloid cells are known producers of certain complement components^22^. Our data indicates that no Factor B was detected in the supernatant of THP-1 cells even post stimulation, despite the increase in C3 levels (**Supplementary Fig. S3C**). Collectively, these data suggest that the increase in Factor B production primarily occurs in intestinal epithelial cells and stromal cells rather than in myeloid cells, in response to certain cytokines that are implicated in gut mucosal immune responses and IBD.

### Gut-derived Factor B protects mice from chemical-induced colitis

To interrogate the role of mucosally-derived Factor B in the gut, we treated global CFB-deficient mice and liver-deficient CFB mice with oral dextran sodium sulfate (DSS). Following DSS treatment, we observed that *Cfb*^−/−^ mice had worse colitis, as measured by weight loss (**Fig. 4A**), shorter colon lengths (an indicator of worse inflammation, **Fig. 4B**), and elevated cytokine and myeloperoxidase (MPO) levels (**Fig. 4C**) compared to *Cfb^f/f^*and *Cfb^f/f^AlbCre*^+/−^ mice. We also observed increased fecal lipocalin-2 (LCN-2) levels in *Cfb*^−/−^ mice compared to *Cfb^f/f^* and *Cfb^f/f^AlbCre*^+/−^ mice (**Fig. 4D**). To assess if the Factor B generated in the colon is functional, we measured AP convertase formation (C3bBbP) in colonic tissue from mice with or without DSS-mediated injury. Our data reveal that C3bBbP levels were increased in the setting of colitis not only in the *Cfb^f/f^* mice, but also in *Cfb^f/f^AlbCre*^+/−^ mice, who have no circulating Factor B (**Fig. 4E**). Pathway analyses from bulk RNASeq on colonic tissues reveal that *Cfb^−/−^* mice had an increased inflammatory response compared to the *Cfb^f/f^AlbCre*^+/−^ mice (**Fig. 4F**). These studies indicate that gut-derived Factor B is active and protects against colitis, independent of circulating Factor B.

**Figure 4.**
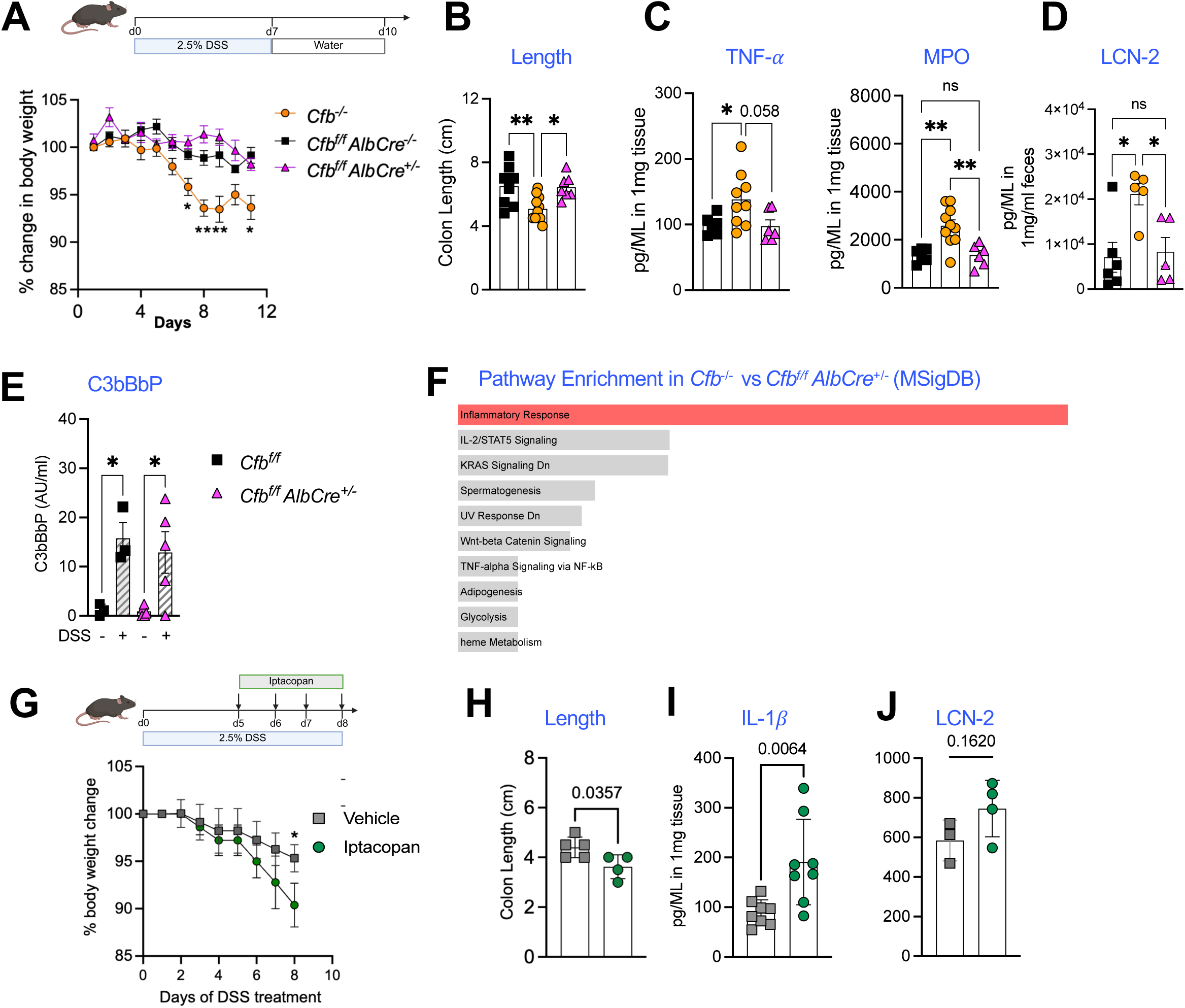
Mucosal Factor B is required for protection in the setting of DSS-induced colitis. (**A**) SPF-housed global knockout (*Cfb^−/−^*), liver-deficient (*Cfb^f/f^ AlbCre^+/−^*) and littermate control (*Cfb^f/f^ AlbCre^−/−^*) mice were administered 2.5% DSS in drinking water for 7 days, followed by conventional water for 3 days. Mice were euthanized on Day 11. Body weights were monitored daily during treatment. (**B**) Colon lengths evaluated at end of study. (**C**) Colonic tissue was homogenized for measuring TNF-α and MPO levels, normalized to weight (per g) of tissue. (**D**) Fecal lipocalin-2 (LCN-2) levels compared at the end of the study. (**E**) Complement activity in the colonic tissue measured using C3bBbP (alternative pathway convertase). (**F**) Differentially regulated pathways upregulated in *Cfb^−/−^* versus *Cfb^f/f^ AlbCre^+/−^* based on bulk RNA-seq of colonic tissue at end of treatment. Pathway analysis conducted using Enrichr (MSigDB). (**G**) Adult C57BL/6 mice housed in SPF conditions were given 2.5% DSS in drinking water for a week. Starting day 5, mice were treated with a daily oral gavage of a Factor B inhibitor, iptacopan (20 mg/kg), or vehicle control. Body weights were monitored daily. (**H**) Colon lengths compared at end of the experiment. (**I**) IL-1β in colonic tissue lysate and (**J**) LCN-2 in fecal samples. Bar graphs show the mean ± SEM, * P < 0.05; ** P < 0.01, ns, non-significant. Unpaired t test when analyzing a two-group categorical comparison, and 2-way ANOVA adjusted for multiple comparisons when analyzing longitudinal data.

Independent of genetic deletion, treatment of wildtype mice with oral iptacopan (an FDA-approved small-molecule Factor B inhibitor)^23^ resulted in worse disease compared to vehicle-treated mice (**Fig. 4G**). Iptacopan is a first-in-class, cell-permeable, oral drug that functions by interfering with the formation of the AP C3 convertase (i.e., C3b<u>Bb</u> complex)^23^. We observed that mice treated with iptacopan had decreased colon length (**Fig. 4H**), higher levels of IL-1β in the colon tissue lysate (**Fig. 4I**) and increased LCN-2 levels in fecal samples (**Fig. 4J**). These observations indicate that Factor B inhibition may be detrimental during colitis.

### Factor B sourced from structural cells in the gut is required for protection against colitis

To interrogate the sourcing of Factor B in the gut from immune and non-immune cells, we queried a recently published scRNA-seq dataset where C57/BL6 mice were treated with DSS colitis for 6 days and evaluated on day 11 after the start of DSS (acute colitis, AC)^24^. Analysis of the colonic cells from this dataset revealed that while the cellular composition of epithelial and myeloid cells dramatically changed during acute DSS treatment, the composition of fibroblasts remained largely unchanged (**Fig. 5A & B**). Within these cell types, we queried the expression of AP pathway components (**Fig. 5C**) and observed that the average expression of *Cfb* increased in all three cell types – myeloid cells, epithelial cells and fibroblasts in the setting of acute colitis. However, both the magnitude of *Cfb* expression and the percentage of cells expressing *Cfb* increased in the epithelial cell population, despite a dramatic reduction in epithelial cells in the setting of acute colitis (**Fig. 5B & C)**.

**Figure 5.**
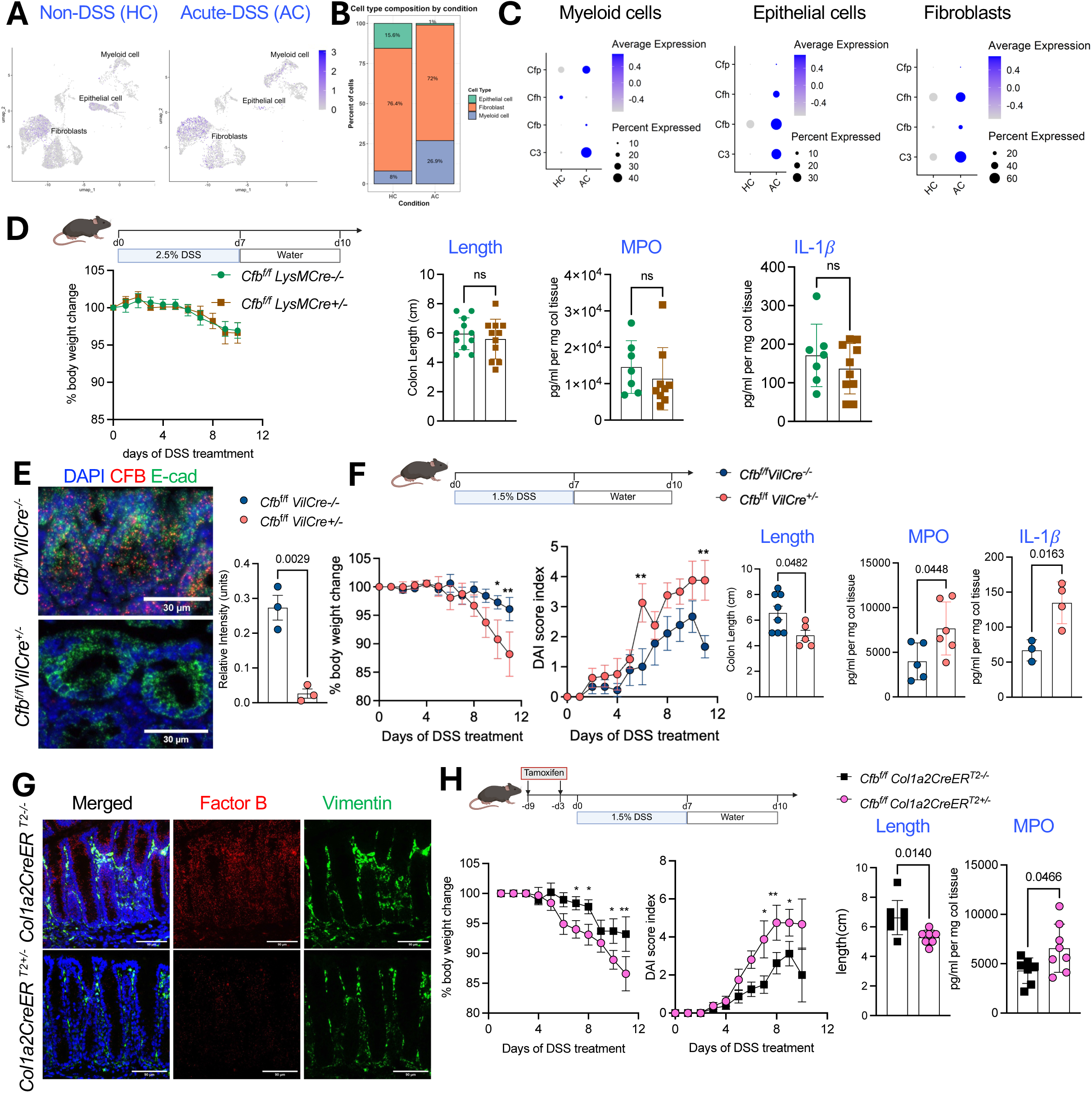
Factor B sourced from intestinal epithelial cells and fibroblasts mitigate the severity of DSS-induced colitis. (**A-C**) Data from scRNA-seq of colonic cells from C57BL/6 mice subjected to acute DSS (acute colitis, AC) was re-analyzed for express of complement genes and compared to mice not treated with DSS (non-DSS, healthy controls (HC), GSE264408). (**A**) UMAP with designated cell clusters showing *Cfb* expression. (**B**) Bar plots depicting changes in the cellular composition of the colonic tissue post-DSS treatment. (**C**) Dot plots showing average gene expression levels of key AP components in the distinct cell clusters of HC (healthy control) mice and those with AC (acute colitis). (**D**) *Cfb^f/f^ LysMCre^+/−^* mice and controls were treated with 7 days of 2.5% DSS in drinking water, followed by 3 days of regular water. Body weights were monitored daily. Colon length was determined at the end of the experiment. Colonic tissue was homogenized for comparing MPO and IL-1β levels, normalized to tissue weight. (**E**) Representative RNAScope images depict staining of colonic tissue from *Cfb^f/f^ VilCre^+/−^* mice and controls. Scale bar =30μm. (**F**) *Cfb^f/f^ VilCre* and controls were treated similarly to (**D**). Body weights and the disease activity index (DAI) were measured daily. Mice were sacrificed on day 11. Colons were harvested to compare the colon length, MPO and IL-1β levels, normalized to weight of tissue. (**G**) *Cfb^f/f^Col1a2CreER^T2+/−^* mice and their controls were treated with tamoxifen every other day for 10 days. Colons were harvested and tissue sections were stained for Factor B (red) and vimentin (green). Scale bar =90*μ*m. (**H**) *Cfb^f/f^Col1a2CreER^T2+/−^* and littermates were subjected to tamoxifen challenge as detailed in (**G**). 3 days after the last dose of tamoxifen, mice were subjected to DSS challenge as detailed in (**D**). Body weights and DAI scores were monitored daily. Colon tissue was collected at day 11 to measure colon length and cytokines in tissue lysate. Data represent 2 independent experiments. Bar graphs show the mean ± SEM, * P < 0.05; ** P < 0.01, ns, non-significant. Unpaired t test when analyzing a two-group categorical comparison, and 2-way ANOVA adjusted for multiple comparisons when analyzing longitudinal data.

Given that Factor B is produced locally in the gut and multiple cell types upregulate it in response to inflammation, we interrogated the necessity and specificity of cell type-intrinsic Factor B production in the host response to colitis. To this end, we first generated *Cfb*^f/f^*LysMCre* mice to interrogate the role of myeloid cell-derived Factor B (**Fig. 5D**). We observed that absence of Factor B in myeloid cells did not influence the course of colitis in our model, as measured by weight loss, colon length, and the disease activity index (DAI), which incorporates measures such as rectal bleeding and fecal consistency (**Fig. 5D**), suggesting that the contribution of Factor B from myeloid cells is redundant in this setting.

To test the contribution of Factor B sourced from gut epithelial cells in the host response to colitis, we generated *Cfb*^f/f^*VilCre* mice. These mice had a constitutively lower co-expression of CFB among E-cadherin-expressing cells (**Fig. 5E**). When *Cfb*^f/f^*VilCre^+/−^*and *Cfb*^f/f^*VilCre^−/−^* littermates were subjected to DSS-induced injury, we observed that the *Cfb*^f/f^*VilCre^+/−^*mice succumbed to disease with increased weight loss, and a worse DAI score compared to the littermate controls (**Fig. 5E**). Additionally, *Cfb*^f/f^*VilCre^+/−^*mice had shorter colon lengths and increased levels of inflammatory cytokines such as MPO and IL-1β compared to the control mice post-DSS (**Fig. 5F**).

Finally, as our single cell analyses revealed that Factor B is also expressed by intestinal stromal cells, we generated *Cfb*^f/f^*Col1a2CreER^T2^* mice with an inducible deletion of Factor B following tamoxifen treatment. We observed that 5 doses of tamoxifen challenge were sufficient to reduce Factor B within colon tissue among *Cfb*^f/f^*Col1a2CreER^T2+/−^*mice compared to littermate controls (**Fig. 5G**). Upon inducing colitis using DSS, we detected greater weight loss and increased disease progression in *Cfb*^f/f^*Col1a2CreER^T2+/−^* mice as compared to *Cfb*^f/f^*Col1a2CreER^T2−/−^* mice (**Fig. 5H**). Additionally, the *Cfb*^f/f^*Col1a2CreER^T2+/−^* mice had worse intestinal inflammation compared to littermate controls following DSS challenge, as shown by reduced colon length and increased MPO levels. These data confirm that Factor B produced by structural cells in the gut mucosa such as epithelial and stromal cells is required for promoting mucosal homeostasis in the setting of colitis.

## DISCUSSION

Factor B has evolved from a primitive complement system present several million years ago, where it was one of the first complement components along with C3-like and MASP-like protein; its original function in invertebrates likely involved opsonization and inflammation^25^. Over time, the Bf/C2f gene duplicated prior to the split of amphibians and mammals, and likely led to the development of the classical, lectin and alternative pathways of complement activation^26^. The evolution of the complement system highlights the necessity of these proteins to protect the lumen of organisms such as protostomes that do not have a liver, and thus, creates a precedent to investigate its role in immune responses at mucosal surfaces, such as the gut.

Both C3 and Factor B facilitate immune responses in humans by attacking microbes to decrease pathogen burden^5, 6^. Their genetic deficiencies are associated with recurrent bacterial infections in both children and adults.^6^ However, their uncontrolled activation has been associated with a multitude of diseases resulting in multisystem involvement, such as atypical hemolytic uremic syndrome, vasculitis, and paroxysmal nocturnal hemoglobinuria^27^. Hence, the clinical interest in complement therapeutics has primary focused on inhibiting complement activation in the circulation^5^. While many of these therapies have been life-changing, most of these trials have not been successful in mitigating diseases involving mucosal surfaces^28, 29^.

There is growing recognition that C3, a central complement component, is locally produced and operative at mucosal sites such as the lung, gut and genitourinary tract^11, 12, 13^.Colonic stromal cells express C3, and intestinal microbes can stimulate local C3 levels^12^. However, little is known about the other complement components, such as Factor B, a key AP component that facilitates amplification of the complement cascade via binding to C3 and its products. Hence, although complement has been implicated in IBD since 1970s^14, 15^, it is unclear as to how it drives disease pathogenesis. Investigators have had challenges in interrogating its contribution to mucosal responses because of the inability to separate circulating complement proteins (primarily made by the liver) from those made locally and are altered in the gut during injury^30^. Additionally, it is unclear if the activation and regulation of these pathways in the gut parallels this balance in the circulation. For example, although we and others have demonstrated the necessity of C3 and Factor B to mitigate cell death^11, 31^, increased C3 and Factor B have also been observed in IBD^14, 15, 16, 17^. Conversely, a loss-of-function genetic variant in *CFB* (*rs4151651*) is strongly associated with IBD severity, indicating an important and dichotomous role of Factor B in IBD, and prompting the need to identify its sourcing and function in gut mucosal responses to injury^7^.

Using both genetic and pharmacological disruption of Factor B activity, we demonstrate that locally sourced Factor B is essential in the host response to colitis. We observe that intestinal epithelial and stromal cells both produce and secrete Factor B, and this sourcing is independent of the circulating, liver-derived Factor B pool. Moreover, Factor B secreted from these cell types lining the lumen increase in response to injury and inflammation. Our data would suggest that this increase in Factor B, a serine protease, would facilitate activation of the luminal AP convertase and cleavage of C3 in the setting of colitis. Correspondingly, we also observe that Factor B expression increases in human tissue specimens during inflammation in IBD. The co-expression of DUOX-2 with Factor B in inflamed tissues suggests local complement activity among epithelial cells in injured regions, potentially as an acute phase response to injury. The necessity of this response is demonstrated using conditional knockouts of structural cell types that form the gut mucosal barrier, which appear to serve as a key source of local Factor B, rather than myeloid cells. Taken together, our data suggest that this core complement component needs to be locally produced as a host defense mechanism to respond to mucosal insults.

Our results challenge certain prior literature which suggest that complement inhibition reduces colitis severity. However, our study highlights the nuances that simply inhibiting the system may not necessarily be beneficial, and the specific complement component, as well as the cell type being targeted may be as, if not more important. For example, most of the prior studies investigating the role of complement in colitis have largely been carried out using whole body knockouts and do not address the effect of locally-derived complement proteins^32, 33^. To our knowledge this is the first in vivo study uncovering the importance of locally sourced Factor B at any mucosal surface, and thus does not negate, but rather builds on these prior observations. Our data indicate that while immune, epithelial and stromal cells contribute to the local pool of Factor B, epithelial and stromal cell-derived Factor B is essential for protection against chemical induced-colitis. Concurrently, key alternative pathway regulators such as CD55 are altered in IBD^34, 35^. Moreover, a biallelic loss-of-function mutation in CD55 results in hyperactivation of complement, angiopathic thrombosis, and protein-losing enteropathy (the CHAPLE syndrome)^36^, which has successfully been treated with pozelimab, a C5-blocking antibody. Thus, understanding how the AP is sourced, activated and regulated in the gut, and how it can be modulated locally to prevent tissue damage is critical for developing targeted therapies^37^.

Our study has certain limitations. First, we deleted Factor B from epithelial and stromal cells based on its single-cell expression from these cell types. Future work will need to address whether the protective effect of cell type-intrinsic Factor B is due to rapid, canonical activation of the complement cascade or its non-canonical/intracellular roles^38^. There is growing recognition that in addition to their role in mounting a rapid innate immune response, complement proteins also play a central role in cellular process such as survival, proliferation and metabolism^38, 39^. Second, we focused on the role of Factor B in an acute colitis model using DSS. The effect of Factor B may be different based on the stressor and the course of disease. We acknowledge that DSS does not correlate to actual disease in humans, but this remains the most suitable model of gut epithelial injury and is currently the most well-established in vivo model to study UC. Future studies would involve other models of injury, such as infection and radiation-induced colitis. Third, we use iptacopan, an FDA-approved Factor B inhibitor as it can be given orally, rather than parenterally. However, we cannot exclude the effects of intracellular inhibition of Factor B, as has been previously shown by other investigators^40^. Our future work would involve separating how best to ameliorate AP activity while preserving the mucosal homeostatic role of Factor B in colitis.

In summary, we show that independent of circulating Factor B, intestinal cell-derived Factor B is required for an appropriate host response to colitis. This protection appears to be derived through Factor B sourced from both epithelial and stromal cells. Our observation that Factor B expression in increased in patients with IBD who have active inflammation also indicates that there is imminent need to better understand and dissect the contribution of local complement proteins and the cell type-specific contribution of these proteins. Future studies should be aimed towards developing targeted approaches to regulate tissue-specific complement activation and comparing them to systemic complement inhibition in order to promote mucosal homeostasis.

## METHODS

### Human data

Sequencing data were obtained from published studies^19, 20^. The description and collection of samples, the data preprocessing is explained in these studies. Deidentified human histology specimens were obtained from Translational Pathology Core Laboratory (TPCL) at UCLA.

### Animal studies

All animal studies were conducted on protocols approved by the Institutional Animal Care and Use Committee at Washington University School of Medicine (WUSM) and the Animal Research Committee at University of California Los Angeles (UCLA). All data are reported from controlled laboratory experiments. Mice ages 8-12 weeks of age on C57BL/6 background were used for all the experiments. All mice were fed sterilized food and water *ad libitum* or experimental water as described below. All animals were maintained in a specific pathogen free (SPF) mouse facility. Commercially available mice were purchased from Jackson Laboratories, Bar Harbor, ME, which included C57BL/6J mice (B6, #006664), B6.Cg-*Speer6-ps1Tg(Alb-cre)21Mgn*/J (*Alb-Cre*, #003574), B6.129P2-Lyz2tm1(cre)Ifo/J (LysM-Cre. #004781), B6.Cg-Tg(Vil1-cre)997Gum/J (*Vil-Cre,* #004586), and B6.Cg-Tg(Col1a2-cre/ERT,-ALPP)7Cpd/2J (Col1a2-CreER, #029567).

*Cfb^−/−^* mice were generated using a CRISPR-Cas9 approach, confirmed and tested for off-target effects by whole genome sequencing (**Supplementary Fig. S2A**). *Cfb*^f/f^ mice were generated by homologous recombination in embryonic stem cells. For this purpose, a targeting vector containing regions homologous to the genomic *Cfb* sequences was constructed (**Supplementary Fig. S2B**). This strategy results in a conditional deletion of 1.4kb of coding sequences encoding for the von Willebrand factor type A (vWFA) domain.

The *Cfb*^f/f^*AlbCre^+/−^*, *Cfb*^f/f^*LysMCre^+/−^*and *Cfb*^f/f^*VilCre^+/−^* were constitutive conditional knockouts, generated by breeding the *Cfb*^f/f^ mice with the respective Cre driver detailed above. *Cre*-negative littermates served as controls for these mice. To generate an inducible deletion of Factor B from Col1a2Cre-expressing fibroblasts, *Cfb*^f/f^ mice were bred with the *Col1a2CreER* strain (Strain #:029567) and were referred to as *Cfb*^f/f^ *Col1a2CreER*^T2^ mice. To delete *Cfb* from fibroblasts, *Cfb*^f/f^*Col1a2CreER^T2+/−^*mice and *Cfb*^f/f^*Col1a2CreER^T2−/−^* mice (referred to as *Cfb*^f/f^ mice in the relevant experiments) were injected with 10 mg/mL, tamoxifen (Sigma-Aldrich, St. Louis, MO, USA) dissolved in sunflower seed oil with 20% ethanol (Sigma-Aldrich) intraperitoneally, every other day for a total of five doses. Three days after the final injection, mice were used for the subsequent experiments.

Germ free (GF) were obtained from the UCLA Goodman-Luskin Microbiome Center Gnotobiotic Core. GF mice were maintained in flexible-film isolators. Sterility was verified at regular intervals using aerobic cultures, anaerobic cultures, and PCR. Co-housed littermates were used as experimental groups and controls to minimize differences in the gut microbiota. For pharmacologic manipulation, littermates of the same sex were assigned a cage, which contained animals of all treatment groups.

### Genotyping

Mouse genotyping was performed by Transnetyx Inc. on tail biopsies done of mice aged 7-14 days old, utilizing duplicate sample processing (real-time TaqMan PCR).

### Sex as a biological variable

Our study investigated data from both male and female human participants, and similar findings are reported for both sexes. Our murine model also used both male and female mice, and no significant differences were observed when analyzed separately.

### Chemical-induced colitis

Mice were administered dextran-sodium sulfate (DSS. 36,000-50,000 molecular weight, Colitis Grade, MP Biomedicals, #9011-18-1) in drinking water ad libitum for seven days, and then subsequently put on regular drinking water form day 7 to day 10. Weight loss, stool consistency and rectal bleeding severity were monitored daily to graph the Disease Activity Index (DAI) and overall disease severity. DAI was scored using the following parameters^41^:

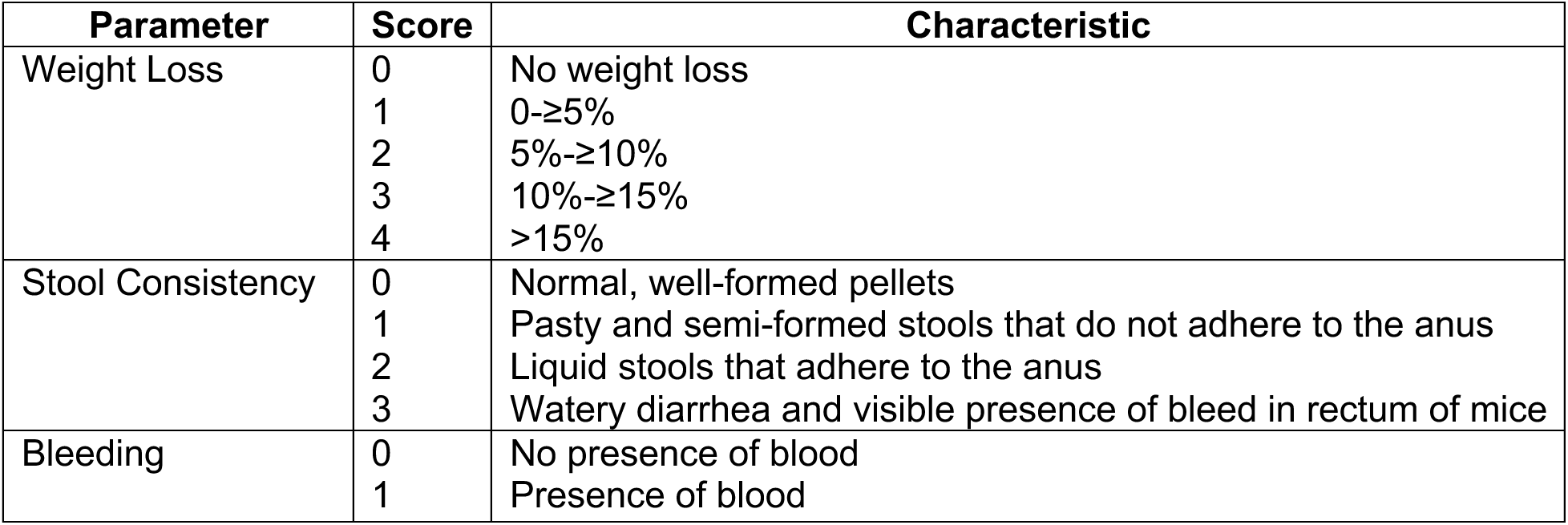

### Pharmacological Factor B inhibition

C57BL/6 mice maintained under SPF conditions were given 2.5% DSS for 8 days. Starting on day 5, mice received once-daily oral gavage of iptacopan (Medchemexpress LLC #HY-127105) at a dose of 20 mg/kg dissolved in 10% DMSO and 90% corn oil. The control mice received oral gavage will vehicle.

### ELISAs

Colonic tissues were weighed and homogenized in T-PER™ Tissue Protein Extraction Reagent supplemented with protease inhibitor at a ratio of 100 mg tissue per 500 µL buffer. Tissue lysates were centrifuged for 10,000g for 30 min at 4 °C. Supernatants were collected and stored at −20 °C. TNF-α (R&D Systems #DY410-05), MPO (R&D Systems DY3667) and IL-1β (DY401) levels were measured by following the manufacturer’s instructions. Pierce™ BCA Protein Assay Kit (Thermo Scientific, #23225) was used to calculate the total protein levels in each lysate though a BCA assay. ELISA results were normalized to the BCA results to account for relative protein levels in each tissue lysate. For the fecal LCN-2 ELISA (R&D ELISA kit #DY1857), fecal samples (frozen) were weighed and resuspended in 0.1% PBS-Tween at a ratio of 100 mg feces per 1 mL buffer. Samples were briefly warmed and vortexed to complete dissolution the fecal material followed by centrifugation for 10 min at 12,000 rpm and 4°C. The supernatants were collected for analysis. AP convertase levels were quantified using a C3bBbP ELISA kit as per manufacturer instructions.

### Immunofluorescence

Formalin-fixed, paraffin embedded sections were first deparaffinized through xylene and ethanol gradients before pressure cooking in citrate-based antigen retrieval solution (Vector Labs., Cat. H-3300) for 30 min. The sections were then blocked with 3% Bovine Albumen (Thermo Fisher, #J10857.22) diluted in phosphate buffered saline (PBS) at room temperature for 1 h. For primary incubation the sections were stained with a 1:50 dilution of polyclonal rabbit anti-goat Factor B antibody (Quidel Ortho, # A311) and a 1:100 dilution of mouse anti-E-Cadherin (BD Sciences, Cat. 610181) overnight at 4°C. Dilutions were done in 3% bovine serum albumin. For secondary immunofluorescent visualization, the sections were stained for 1 h at room temperature with a 1:200 dilution of donkey anti-goat Alexa Fluor 555 (Thermo Fisher, # A21432) for Factor B and a 1:250 dilution of donkey anti-mouse Alexa Fluor 647 (Thermo Fisher, #A31573) for E-Cadherin. The nucleus was then cross stained and mounted with Diamond Antifade mountant with DAPI (Thermo Fisher, #P36962). Images were acquired on ECHO Revolve Rvl2, a widefield microscope (ECHO, San Diego).

### RNAscope

Multiplex fluorescent RNAscope assays were preformed using the RNAscope Multiplex Fluorescent V2 Assay kit (ACDBio, Cat. 323100). The RNAscope target probes used were *Cfb* (ACDBio, Cat. 317051), *Vil1* (ACDBio, Cat. 463301-C3), and *Col1a1* (ACDBio, Cat. 319371-C4). Formalin-fixed, paraffin embedded sections were first deparaffinized through xylene and ethanol gradients before sitting in hydrogen peroxide for 10 minutes at room temperature.

The sections were then placed in a boiling target retrieval solution (ACDBio, Cat. 322000) for 15 minutes at >99°C. After target retrieval, sections were rinsed in distilled water for 15 seconds and dehydrated with 100% ethanol for 3 minutes. Protease Plus reagent (ACDBio, Cat. 322331) was added on each section for 30 minutes at 40°C, then washed off twice in distilled water. Sections underwent probe hybridization and signal amplification according to ACDBio protocols. Images were taken the next day on ECHO Revolve Rvl2, a widefield microscope.

Fluorescence was quantified using FIJI ImageJ version1.54p analysis. For each image, the images were uploaded to ImageJ and regions of interest (ROI) were traced through free-hand selection and added to the ROI Manager. Additionally, a section of background was selected and added to the ROI Manager. The image channels were then split, and fluorescence was measured for each channel using the previously set ROIs in the ROI Manager. Once the raw integrated density data was obtained, background correction was applied by using this equation: Corrected Integrated Density = Raw Integrated Data -(Background Mean x Area of ROI). Finally, Factor B fluorescence is divided by E-Cadherin fluorescence to show the amount of Factor B present per E-Cadherin stain.

### In vitro experiments

Caco-2 epithelial cells were maintained in DMEM supplemented with 10% fetal bovine serum (FBS) and 1% Penicillin–Streptomycin (P/S). Primary human fibroblasts were cultured in DMEM/F12 medium containing 10% FBS and 1% P/S. THP-1 human monocytic cells were cultured in RPMI-1640 medium supplemented with 10% FBS and 1% P/S. All cell lines were incubated at 37°C in a 5% CO₂ atmosphere. For all experiments, cells were seeded in 6-well plates at a density of 1.5 × 10⁶ cells per well and allowed to grow until they reached approximately 70–80% confluence. For suspension THP-1 monocytes, cells were allowed to equilibrate after seeding before cytokine treatment. Before cytokine stimulation, Caco-2 and fibroblast cultures were washed once with sterile phosphate-buffered saline (PBS) to remove residual serum, followed by replacement with serum-free DMEM (Caco-2) or serum-free DMEM/F12 (fibroblasts) containing 1% P/S. THP-1 cells were similarly washed with PBS and switched to serum-free RPMI-1640 supplemented with 1% P/S. All cell types were treated with human recombinant IL-1β and TNF-α at a final concentration of 50 ng/mL each. Cytokine stimulation was carried out for 24 hours under standard culture conditions. Following cytokine treatment, culture supernatants were collected from each cell type and centrifuged at 300 × g for 5 minutes to remove cellular debris. The clarified supernatant was then concentrated using a 3 kDa Amicon Ultra centrifugal filter unit according to the manufacturer’s instructions. The concentrated samples were immediately stored at −80°C until use for immunoblotting. For preparation of the lysate, the supernatant was collected, and cells were washed with ice-cold PBS and lysed directly in the wells using RIPA lysis buffer supplemented with protease and phosphatase inhibitors. Lysates were incubated on ice for 15 min to ensure complete protein extraction, then centrifuged at 12,000 × g for 10 min at 4°C. The resulting protein lysate was collected and stored at −80°C.

### Immunoblotting

Protein samples (20 µL) were mixed with 4 µL of 4x loading dye, incubated at 98 °C for 10 min, and loaded onto pre-cast SDS–PAGE gels. Electrophoresis was performed using Tris–Glycine SDS running buffer at 200 V for approximately 40 min until the dye front reached the bottom of the gel. Following electrophoresis, gels were equilibrated in deionized water for 5 min with gentle shaking. Proteins were transferred to nitrocellulose membranes using a pre-assembled iBlot Western blot transfer stack (Thermo Fisher Scientific). The gel was placed on the membrane with all layers assembled according to the manufacturer’s instructions, ensuring that no air bubbles were present. Transfers were performed in the iBlot device at 25 V for 3 min under high cooling conditions. After transfer, membranes were rinsed twice with TBS-T for 5 min, then blocked in 3–7% skim milk (prepared by dissolving 0.75–1.75 g of skim milk powder in 25 mL of TBS-T) for 1 h at room temperature with gentle shaking. Membranes were incubated overnight at 4 °C with a goat polyclonal anti-human/mouse Factor B primary antibody (Complement Technologies #A213) diluted 1:10,000 in 3–7% milk/TBST. Following three 5-minute washes in TBS-T, membranes were incubated for 1 h at room temperature with an HRP-conjugated donkey anti-goat secondary antibody diluted 1:10,000 in 3–7% milk/TBST. After two additional TBST washes, membranes were incubated for 5 min with a 1:1 mixture of chemiluminescent substrate solutions and imaged using a GelDoc imaging system.

### Statistical analyses

Data are presented as scatterplots showing individual data points, and dispersion is shown by mean and standard deviation. Unpaired t test was used for comparing two groups, assuming equal variance, and ordinary one-way analysis of variance (ANOVA) test with multiple comparisons testing was used for more than two groups. A P value of <0.05 was considered significant, although values of >0.05 have been reported when there was evidence of a trend. Prism v10.3.0 (GraphPad) was used for statistical analysis.

### Single-Cell RNA sequencing (scRNA-seq)

scRNA-seq analyses was conducted at the Genome Technology Access Center @ McDonnell Genome Institute at Washington University. Data were processed using the CellRanger pipeline (version 8.0.1) with default settings, aligning reads to the mouse mm10 (GENCODE vM23/Ensembl98) reference genome. The resulting gene-barcode matrix was analyzed using the Seurat package (version 5.1.0) in R (version 4.4.2). Cells with at least 500 detected genes and fewer than 40% mitochondrial reads were retained. Normalization and variance stabilization were performed using the SCTransform method. Dimensionality reduction was conducted using 30 principal components, followed by Louvain clustering and UMAP visualization. Cluster-specific markers were identified using the PrepSCTFindMarkers and FindAllMarkers functions in Seurat, employing the Wilcoxon rank sum test and a log fold change threshold of 0.25. Broad and fine-grained cell types were annotated based on canonical marker genes.

### Bulk RNA sequencing (RNA-seq)

Bulk RNA-seq analyses was conducted at the Genome Technology Access Center @ McDonnell Genome Institute at Washington University and at the Technology Center for Genomics & Bioinformatics (TCGB) at UCLA. Raw paired-end FASTQ files were processed using Partek Flow software. Adapter trimming and quality filtering were performed, followed by alignment to the mouse reference genome (mm10) using the STAR aligner. Gene-level quantification was performed in Partek Flow to generate read count matrices. Downstream analysis was conducted in R (version 4.4.2) using the DESeq2 package (version 1.46.0). The data consisted of paired samples. Lowly expressed genes were filtered out, and count data were normalized using DESeq2’s median-of-ratios method. Principal component analysis (PCA) was used to assess sample variation. Differential expression analysis was performed using DESeq2, with p-values adjusted using the Benjamini–Hochberg method. Genes with an adjusted p-value < 0.05 and absolute log2 fold change ≥ 1 were considered significantly differentially expressed. For gene set enrichment analyses and pathway analyses, the EnrichR web-based tool and GSEA algorithm were used. Ensembl gene IDs were converted to gene symbols using the org.Mm.eg.db annotation package. Visualization of results, including PCA plots, heatmaps, and volcano plots, was performed using ggplot2 and pheatmap packages in R.

### Spatial transcriptomic analysis

Visium data were obtained from the GEO (GSE189184)^20^. The dataset included three sample groups — healthy controls (HC), ulcerative colitis (UC), and checkpoint inhibitor-induced colitis (CC). For the purposes of this study, only the HC and UC samples were included in downstream analyses. All analyses were performed using the normalized Visium data as provided, without applying additional preprocessing or quality control steps beyond those described in the original study. Average normalized expression of *CFB* was compared between HC and UC samples, and co-expression of *CFB* and *DUOX2* was assessed among the cells at group-level. Spatial expression patterns for *CFB*, as well as co-localization of *CFB* and *DUOX2* across Visium spots, were visualized using Loupe Browser (10x Genomics). Similarly, C3 expression across Visium spots were also visualized.

### Study approval

Animal experiments were conducted in accordance with an approved protocol at University of California-Los Angeles (ARC-2024-060, ARC-2024-101, ARC-2004-056) and at Washington University School of Medicine (23-0374).

### Data availability

The deidentified data has been deposited on GEO and will be made publicly available at the time of publication.

## AUTHOR CONTRIBUTIONS

Designing research studies: AT, NM, SN, HWK, KM, MC, DHK, HSK

Conducting experiments: AT, NM, SN, KT, BY, JH, JMS, CSS

Acquiring data: AT, NM, SN, KT, BY, JH, JMS, DHK

Analyzing data: AT, AN, NM, SN, KT, JH, JMS, KM, DHK, HSK

Writing the manuscript: AT, AN, NM, SN, KM, DHK, HSK

Revising the manuscript: HWK, ZL, MAC, DHK, HSK

Funding acquisition: HWK, MAC, DHK, HSK

*Co-first author contributions: AT led work on the experimental models while AN led the work on data analysis.

## SOURCES OF SUPPORT

**H.S.K**.: NIH (R01HL166449), WashU Medicine-Rheumatic Diseases Research Resource-based Center (RDRRC, P30AR073752), and Jonsson Comprehensive Cancer Center Seed Grant

**D.H.K**.: NIH (K01DK133670), The UCLA Goodman-Luskin Microbiome Center, Lawrence C. Pakula, MD IBD Education and Innovation Fund (Washington University)

**H. W. K:** NIH (R01DK128142)

**M. A. C**: NIH (R01AI167285) and (R01CA278197)

