## Supplementary Figures for "Mucosally sourced complement factor B modulates the host response to colitis"

### Supplementary Figure S1

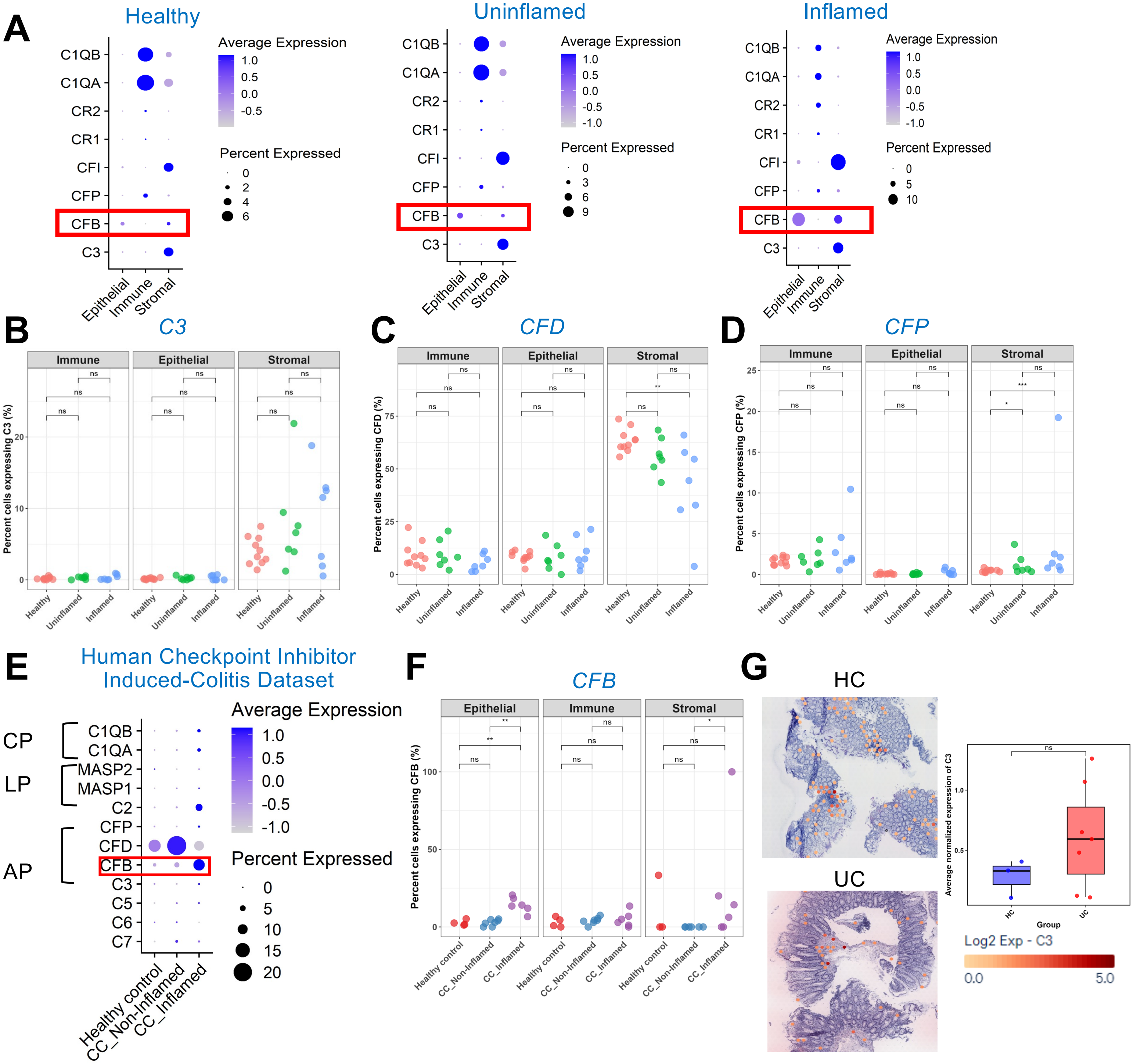


**Supplementary Figure S1. Complement expression in tissue specimens from patients with IBD.** (**A**) Dot plot illustrating the expression of complement genes in intestinal tissue biopsies from healthy controls and from inflamed and uninflamed tissues of patients with ulcerative colitis (UC) distributed over epithelial, stromal and immune cell types. Original data queried from Single Cell Portal: SCP259. (**B-D**) Plots depict expression of (**B**) *C3*, (**C**) Factor D (*CFD*), and (**D**) properdin (*CFP*) across denoted colonic cell types from intestinal tissue biopsies of healthy controls and patients with UC from the same dataset. Each dot depicts individual donor samples. * P <0.05, *** P < 0.001, ns not significant, Wilcoxon rank test. (**E**) Dot plot illustrating the expression of complement genes representing the classical (CP), lectin (LP) and alternative (AP) pathway in intestinal tissue biopsies from uninflamed and non-inflamed regions of patients with checkpoint inhibitor-induced colitis (CC, GSE189184). (**F**) Plots depict *CFB* expression across denoted colonic cell types in the same dataset from intestinal tissue of patients with CC (non-inflamed vs inflamed regions) and healthy controls. * P <0.05, *** P <0.001, ns, not significant, Wilcoxon rank test. (**G**) Representative spatial transcriptomic slides (Visium) showing *C3* expression in intestinal tissues obtained from patients with UC compared to healthy controls (HC, GSE189184). Graph depicts quantification of *C3* expression from 3 HC and 7 UC specimens. Error bars represent standard error of the mean (SEM). ns (non-significant), Mann-Whitney U test.

### Supplementary Figure S2

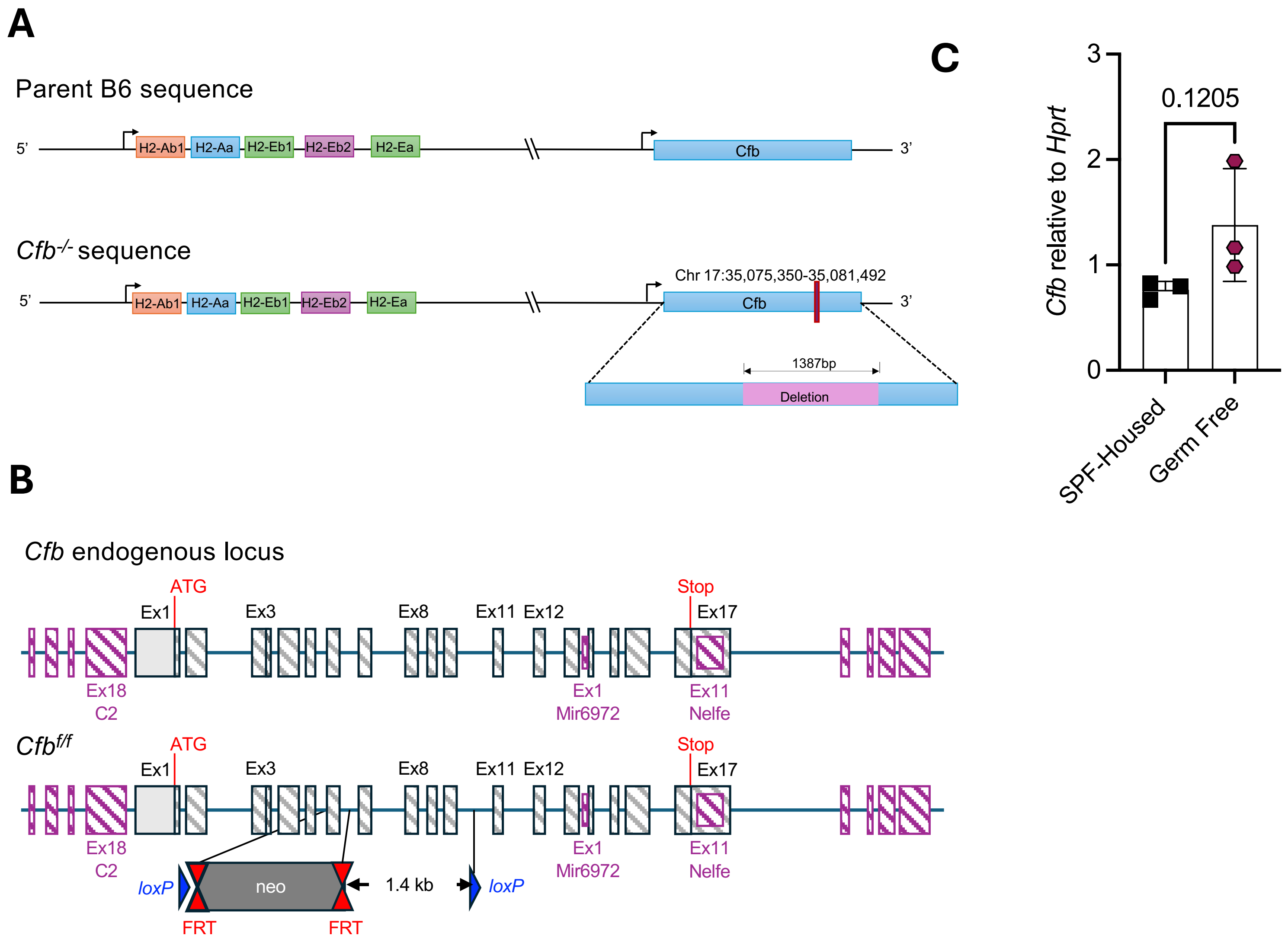


**Supplementary Figure S2. Transgenic mice utilized for Factor B deletion.**
**(A)** Transgenic *Cfb*$^{-/-}$ mice were generated by introducing a 1287bp deletion in the *Cfb* gene on chromosome 17. Targeted embryonic stem (ES) cell clones containing the appropriate genetic changes that ablate expression of the gene of interest are transferred to the blastocoele cavities of 3.5-day blastocyst embryos (generally 10-15 ES cells/host embryo) and, in turn, the embryos are transferred to surrogate mothers where gestation is completed. (**B**) *Cfb*$^{f/f}$ mice were generated by homologous recombination in ES cells. Diagram not depicted to scale. Hatched rectangles represent *Cfb* coding sequences, grey rectangles indicate non-coding exon portions, solid lines represent chromosome sequences, overlapping and adjacent genes are represented as purple rectangles. The neomycin positive selection cassette is indicated. loxP sites are represented by blue triangles and FRT sites by double red triangles. The initiation (ATG) and Stop (stop) codons are indicated. The size of the flanked *Cfb* sequence that would be conditionally deleted is specified. (**C**) *Cfb* expression in the colon tissue from mice housed in a germ-free facility, compared to those housed in specific pathogen-free (SPF) conditions. Values normalized to *Hprt1.*

### Supplementary Figure S3

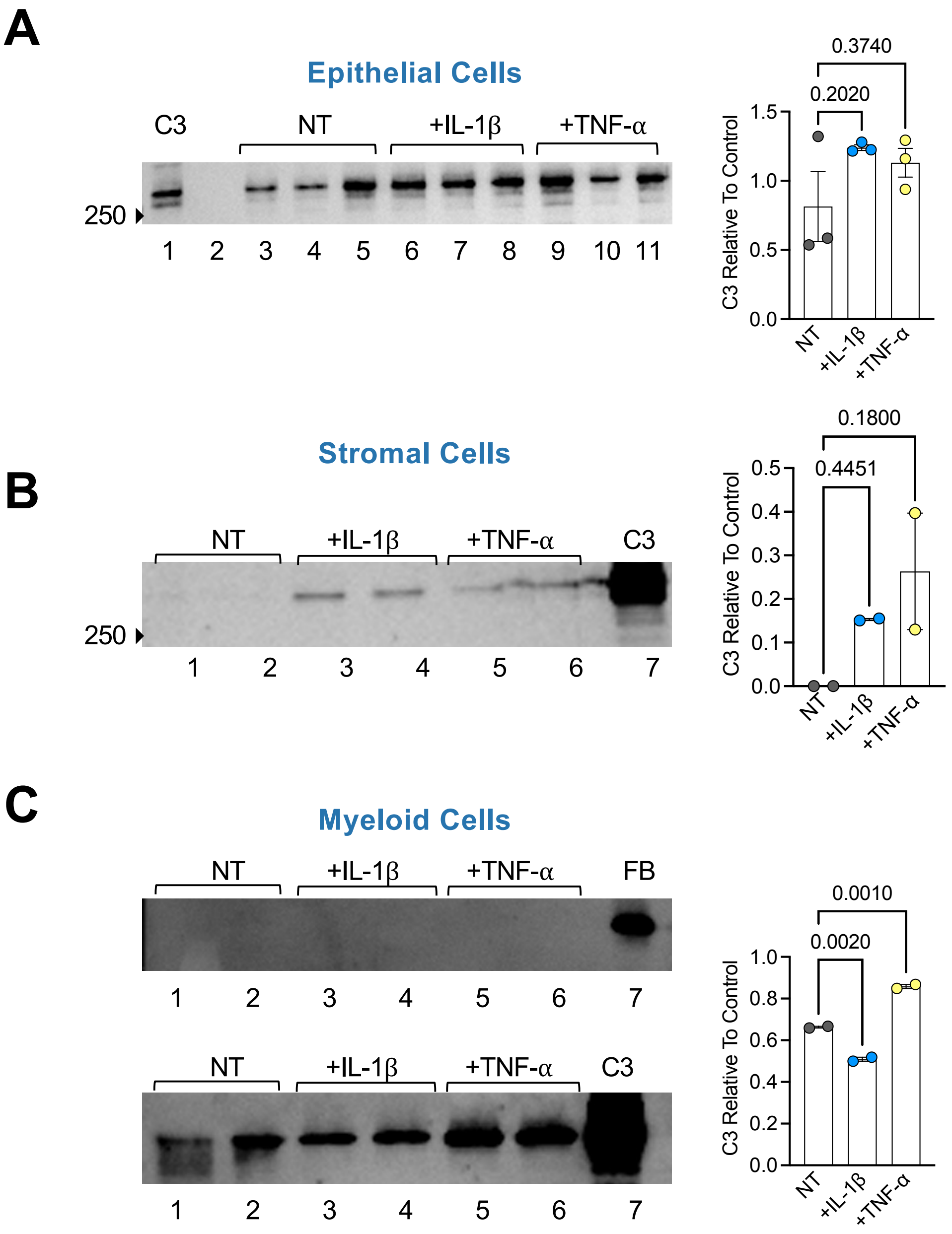


**Supplementary Figure S3. C3 secretion from colonic cells following cytokine stimulation.**
(**A**) Human epithelial Caco-2 cells were incubated in serum-free media alone (non-treated, NT), media with rIL-1β (50ng/ml) or TNF-α (50ng/ml). Supernatant was collected at 24 h, concentrated 50x, and analyzed by immunoblotting. Graph depicts C3 protein quantification relative to the loaded control. (**B**) Human primary fibroblasts were incubated in serum-free media alone (NT), media with rIL-1β (50ng/ml) or TNF-α (50ng/ml). Immunoblot depicts C3 in supernatant at the 24 h time point. Graph depicts C3 protein quantification relative to the loaded control. (**C**) Cells from the human monocytic line THP-1 were cultured and stimulated with indicated cytokines for 24 h, similar to (B) and (C), and their supernatant was probed for Factor B (upper panel) and C3 (lower panel). For all 3 panels, the same volume of supernatant was loaded in each well. Each dot represents an individual replicate. Bar graphs show the mean ± SEM. All immunoblots were repeated at least twice. Statistics performed using an ordinary one-way ANOVA test adjusted for multiple comparisons.
